# Intrinsic neural diversity quenches the dynamic volatility of neural networks

**DOI:** 10.1101/2022.08.25.505270

**Authors:** Axel Hutt, Scott Rich, Taufik A Valiante, Jérémie Lefebvre

## Abstract

Heterogeneity is the norm in biology. The brain is no different: neuronal cell-types are myriad, reflected through their cellular morphology, type, excitability, connectivity motifs and ion channel distributions. While this biophysical diversity enriches neural systems’ dynamical repertoire, it remains challenging to reconcile with the robustness and persistence of brain function over time. To better understand the relationship between heterogeneity and resilience, we analyzed both analytically and numerically a non-linear sparse neural network with balanced excitatory and inhibitory connections evolving over long time scales. We examined how neural diversity expressed as excitability heterogeneity in this network influences its dynamic volatility (i.e., its susceptibility to critical transitions). We exposed this network to slowly-varying modulatory fluctuations, continuously interrogating its stability and resilience. Our results show that excitability heterogeneity implements a homeostatic control mechanism tuning network stability in a context-dependent way. Such diversity was also found to enhance network resilience, quenching the volatility of its dynamics, effectively making the system independent of changes in many control parameters, such as population size, connection probability, strength and variability of synaptic weights as well as modulatory drive. Taken together, these results highlight the fundamental role played by cell-type heterogeneity in the robustness of brain function in the face of change.

**Significance Statement:** Contemporary research has identified widespread cell-to-cell intrinsic diversity in the brain, manifest through variations in biophysical features such as neuronal excitability. A natural question that arises from this phenomenon is what functional role, if any, this heterogeneity might serve. Combining computational and mathematical techniques, this interdisciplinary research shows that intrinsic cell-to-cell diversity, far from mere developmental noise, represents a homeostatic control mechanism, promoting the resilience of neuronal circuits. These results highlight the importance of diversity in the robustness and persistence of brain function over time and in the face of change.

## 1 INTRODUCTION

Neural systems exhibit surprisingly reliable behavior across a lifespan. Despite high phenotypic variability [1–4], learning related plasticity changes [5], and constant alterations in neuromodulatory tone [6–11] and circuit topology [7, 12], neural dynamics remain qualitatively invariant in healthy brains over extended time scales. This is a signature of the brain’s manifest resilience, where its dynamics persist despite changes in intrinsic and/or extrinsic control parameters, preserving associated function [13–17]. In contrast, the failure to regulate such perturbations may predispose neural systems to dynamic volatility: qualitatively distinct dynamics following changes in stability, resulting from critical transitions [18]. Such volatile dynamics in neural systems often arise from disease states: for example, changes associated with epilepsy [19], stimuli [20], or modulatory fluctuations associated with circadian and/or multidien rhythms [21] may cause these systems to slip towards critical transitions, such as recurrent seizures [22–24].

The resilience of neural circuits has been thoroughly studied through pioneering experiments in the crab and lobster stomatogastric ganglia (STG) network [1, 25]. These experiments revealed highly stable, robust and invariant rhythmic activity despite pervasive phenotype heterogeneity, even when exposed to severe environmental perturbations [1]. These discoveries in neuroscience echo a long history, primarily in the field of macroecology, of experimental and theoretical studies examining the relationship between biodiversity, stability, and the resilience of ecosystems and food webs over time (see [14, 16, 26–32] and references therein), which typify the well known ‘stability-diversity’ debate [14, 16, 30, 33]. In this setting, resilience is a system’s propensity for invariance and ability to retain its (in)stability in response to changing control parameters. In contrast, volatile systems are associated with changes in stability and critical transitions, also called bifurcations [13,15,18,30]. A confluence of theoretical studies in macroecology have explored this question and shown (see [33] and references therein) that diversity often renders a system volatile. Combined graph-theoretic and spectral approaches have shown that complex networks tend to lose stability when population sizes increase [14, 32, 34], coupling weights are too strong and/or diverse [14, 26–28, 31, 35], connection probability is too dense [14, 31, 34–36], or when connectivity motifs become too heterogeneous [37].

These questions have been examined by neuroscientists as well: numerous experimental [1, 4, 25, 38–43] and theoretical [43–50] studies have explored the influence of cellular heterogeneity, seemingly the norm in the brain [51–56], on neural dynamics and communication. Furthermore in the context of disease states, excitability heterogeneity can stabilize neural dynamics away from pathological brain states [43]. Collectively, these studies have shown that cell-to-cell diversity stabilizes “healthy” dynamics to optimize responses, learning, information flow and coding capacity by tuning neural networks towards criticality [57], a regime that balances quiescence and excitability while residing between stability and instability. Despite these advances, linking single neuron attributes with emergent physiological activity that undergirds the persistence of brain function remains inaccessible by current experimental techniques.

Inspired by decades of theoretical work in macroecology, we extended spectral theory of random networks [34, 58, 59] and applied it to neuroscience to study the impact of phenotype diversity on the brain’s resilience over extended time scales. We considered a generic large-scale non-linear neural network with sparse balanced excitatory and inhibitory connections, over time scales spanning minutes, hours and/or days to examine the persistence of its dynamics. We exposed this network to a slowly fluctuating modulatory input, a control parameter that is continuously interrogating the system’s stability. Over such time scales, slow modulation influences neural activity in a manner mimicking fluctuations during the resting state resulting from modulatory [6–11], environmental [1, 25, 60], and/or stimuli-induced perturbations, for instance. To quantitatively determine a system’s resilience or volatility, we leveraged spectral theory for large random systems [34,58,59], commonly used in macroecology to examine the stability of complex natural systems, such as food webs [14, 16, 26–32]. Through this framework, we analyzed the statistical properties of eigenvalues resulting from changes in network size, synaptic weights, connectivity motifs, modulatory drive, and cell-to-cell intrinsic diversity amongst neurons. In so doing, we looked beyond the stability of the system to how this stability responds to intrinsic and/or extrinsic changes, in order to understand how excitability heterogeneity predisposes balanced neural systems to stability transitions.

We begin these explorations by showing that excitability heterogeneity, one of many types of intrinsic phenotypic diversity (see Discussion), renders networks less prone to sudden shifts in stability. Excitability heterogeneity refers to cell-to-cell variability in firing rate thresholds (see Methods). We specifically focused on excitability heterogeneity, given that neuronal excitability is a primary mechanism targeted by intrinsic plasticity mechanisms in learning [61], and which is altered in pathological states like epilepsy [43] and neuropsychiatric conditions [62]. We leveraged spectral theory for large random systems to reveal that excitability heterogeneity implements a generic control mechanism promoting: 1) homeostasis, by tuning the distribution of eigenvalues complex plane in a context-dependent way; and 2) resilience, by anchoring this eigenvalue distribution and gradually making it less dependent on modulatory influences. We explored how excitability heterogeneity can influence system resilience to ”insults” like increases in network size, connection probability, strength and variability of synaptic weights, and modulatory fluctuations which promote stability transitions. We found that intrinsic excitability heterogeneity rendered the network more resilient to these insults, a generic feature that was further preserved across a wide range of network topologies. These findings are particularly relevant to learning where synaptic plasticity, unless stabilized by homeostatic mechanisms, would lead to runaway (i.e., unstable) activity [5, 63, 64]. Taken together, these results provide new vistas on the role of a fundamental organizing principle of the brain - neural diversity [51–53] - in brain resilience.

## 2 RESULTS

Neural systems display activity that remains qualitatively invariant over extended time scales highlighting their resilience in the face of changes in connectivity, development and ageing, pathological insults, and exposure to perturbations such as stimuli and/or modulatory influences [6–11]. To better understand the mechanisms underlying such resilience and how it is influenced by excitability heterogeneity, we developed a mathematical framework in which long term stability can be analytically quantified (see MATERIALS AND METHODS). We built and analyzed a large-scale, balanced and sparse network with excitatory and inhibitory connections (see Fig. 1) whose dynamics extend over time scales spanning minutes, hours and/or days. This model is both flexible and general, encompassing a wide range of population-based models involving excitatory and inhibitory interactions. It relates network size, the mean activity of neurons, their mutual synaptic connectivity, their individual level of excitability, and the influence of slowly varying modulatory inputs. We required that neurons were exposed to balanced synaptic connectivity such as seen experimentally [65, 66], in which the net sum of excitatory and inhibitory synaptic weights is zero. We further selected connection probabilities reflecting those observed experimentally [67].

**Figure 1.**
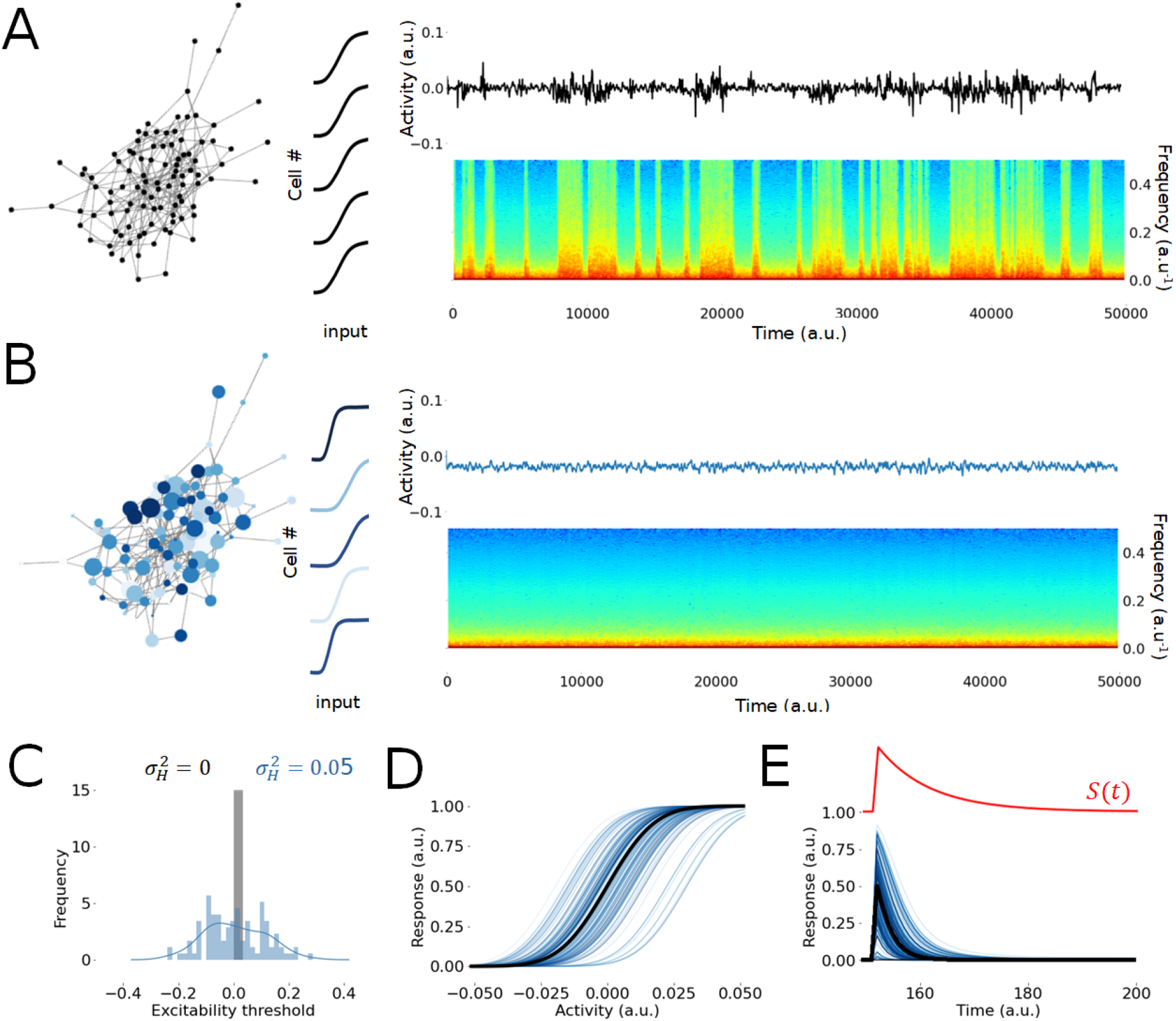
Intrinsic neural diversity promotes the resilience of balanced networks. **A**. Homogeneous networks are composed of neurons with the same biophysical properties, yielding identical excitability profiles (left). Mean network activity (see MATERIALS AND METHODS) displays recurring sudden shifts in stability (i.e. bifurcations) characterized by transitions between states of low- and high-frequency activity. **B** Intrinsic excitability heterogeneity results in variability in the excitability profile of neurons (left) while suppressing shifts in stability. Low-frequency activity persists. **C**. The distribution of excitability thresholds in homogeneous networks (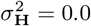; grey histogram) displays zero variance, while intrinsic diversity in heterogeneous networks increases excitability threshold variability (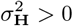; blue histogram). Thresholds were sampled from a normal distribution of mean zero and variance 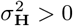 (see main text). **D** Such heterogeneity is reflected in the firing rate response functions which encapsulates the excitability profile of each neuron. In the homogeneous case (black lines; indistinguishable from each other), response functions are identical, but differ in presence of heterogeneity (blue lines). **E** Homogeneous neurons exhibit both the same baseline activity and response to perturbations (black lines; indistinguishable from each other). In contrast, heterogeneous networks exhibiting diversity in excitability yield diversified baseline activities and responses to perturbations (blue lines). The perturbation applied (i.e. *S*(*t*)) is plotted in red (top). The input applied is a filtered step function i.e. 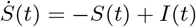 with *I*(*t*) = 0.05 at *t* = 150 and *I*(*t*) = 0 otherwise. Other parameters are given by *d* = −1, *β* = 15 and 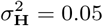.

Within this framework, we can tune the intrinsic excitability of each individual neuron, resulting in increasingly heterogeneous networks; without such variability, the network remain homogeneous. It is well-known that balanced networks are prone to volatility, i.e., susceptible to stability transitions [68, 69]. To confirm this, neurons in the network were collectively exposed to a random, slowly varying modulatory input, mimicking excitability changes in neural activity arising from endogenous and/or exogenenous control parameter changes (i.e., neuromoduation, temperature, etc.) [6–11]. Such a slowly varying modulatory input continuously interrogates network stability and therefore is an ideal tool to expose the system’s resilience. As expected from this context, the homogeneous network (Fig. 1A) was predisposed to volatility through recurring changes in stability. Frequent sharp transitions between states of low- and high-frequency dynamics could be observed in the network’s mean activity (see MATERIALS AND METHODS), and confirmed using power spectral analysis. Such transitions index states of instability, characterized by elevated high-frequency activity, and are reminiscent of dynamics seen in electrophysiological recordings during seizures [70].

However, heterogeneous networks did not exhibit such transitions in response to an identical modulatory input (Fig. 1B). Instead, intrinsic excitability variability was found to suppress these transitions, and low-frequency activity persisted throughout. Intrinsic variability amongst neurons was implemented by varying the effective firing rate response functions, reflecting diverse degrees of cellular excitability. We randomized firing rate response thresholds in which excitability is sampled from a normal distribution of mean 0 and variance 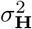 (Fig. 1C; see MATERIALS AND METHODS). This way of characterizing heterogeneity is well aligned with experimental evidence that excitability is a key target for intrinsic plasticity mechanism [3, 43, 61, 62, 71], and echoes numerous previous studies on heterogeneous networks [43, 47, 49, 50, 72]. This heterogeneity leads to differing response functions for the individual neurons (Fig. 1D) as well as variable responses to perturbations like those ocurring due to fluctuations in modulatory input (Fig. 1 E).

### 2.1 Resilience to modulation across time scales

We further characterized the dynamic volatility of our homogeneous network in Fig. 2A. For the parameters chosen, neuronal activity was characterized by alternating epochs of stability and instability, as portrayed in Fig. 1A. This volatility was further confirmed by numerically computing Lyapunov exponents, which we found repeatedly changed sign (Fig 2B), as expected from the theory of nonlinear balanced networks [68, 69]. Slowly driven by the modulatory input, the network dynamics displayed seemingly stable behavior over short time scales. The activity of individual neurons appears smoothly driven by the modulatory input. Such stable dynamics indexes states in which neural activity is stable, and relaxes back to equilibrium after perturbations; at these temporal scales, this corresponds to asynchronous neural firing. We emphasize that fast noise-like fluctuations, commonly present and expected in neural recordings, are here absent, as a consequence of the slow time scale considered. In contrast, other periods were characterized by unstable neural activity (Fig 2C) in which the activity of individual neurons diverge away from equilibrium. Such periods of instability result from modulation-driven critical transitions [18] in which neural activity departs from stability, and may diverge, become synchronous and/or chaotic. To confirm the robustness of these results, we computed the mean Lyapunov exponent across independent realizations of the network connectivity and independent trials, in which the system possesses the same parameters (i.e. connection probability, synaptic weights, proportions of excitatory and inhibitory couplings) but exhibit different configurations and exposed to variable modulatory input. As shown in Fig. 2D, persistent positive mean Lyapunov exponent with large variance could be observed, confirming volatility. Collectively, these observations show that slowly fluctuating modulatory input may expose the volatility of homogeneous networks by revealing sudden stability transitions and dynamical regimes that are qualitatively distinct. This exemplifies non-resilient behavior.

**Figure 2.**
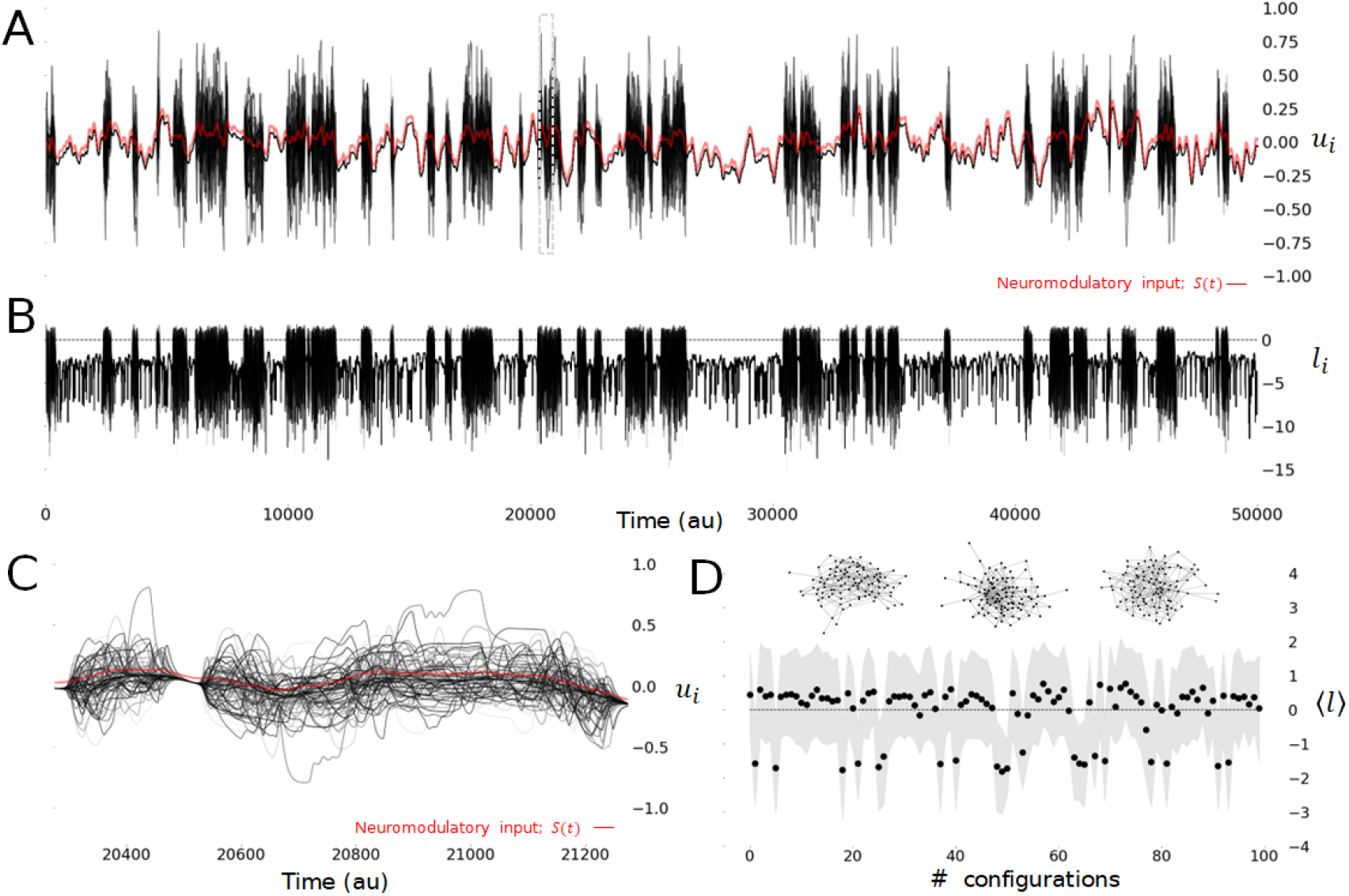
Dynamic volatility of homogeneous networks when exposed to modulatory input over long time scales. **A** Dynamics of a homogeneous network where all neurons possess the same level of excitability. Averaged neuron activity over an extended time scale. A slowly varying modulatory input (*S*(*t*), red line) continuously interrogates the network stability, leading the network through alternating epochs of stability and instability which typifies high volatility. Individual nodes (*u_i_*; black curves) display transient unstable dynamics, alternating with periods of stability. **B** Lyapunov exponents (*l_i_*) – computed numerically based on time series of each neuron across time - delineate periods of stability and instability. The network is unstable whenever Lyapunov exponents are positive, and stable otherwise. This behavior exemplifies volatile (frequently changed) stability. **C**. For some epochs, the network is stable and exhibits dynamics smoothly driven by the modulatory input. Such dynamics corresponds to regimes in which neural activity relaxes back to the equilibrium after a perturbation. These stable periods alternate with epochs of instability in which the activity of the neurons diverge: Such dynamics are characterized by diverging, synchronous and/or chaotic neural activity, and do not relax back to the equilibrium after a perturbation. Fast noise-like fluctuations commonly observed in neural signals are here absent, being averaged out at these slow time scales. **D** Network mean Lyapunov exponent 〈*l*〉 over independent network configurations, realized using identical parameters but different net connectivity. Fluctuations overlapping the horizontal line (〈*l*〉 = 0; black dashed line) indicate volatility. Gray shading indicates ± SD computed over independent trials of duration *T* = 3000a.u.. Other parameters are *N* = 100, *ρ* = 0.05, *f* = 0.8, *μ_e_* = 0.08, *β* = 50, 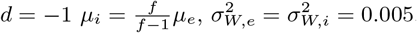, *B* = −0.05.

### 2.2 Dynamic volatility of homogeneous networks

To better understand the dynamics observed in Fig 2 and the underlying mechanisms hindering the resilience of homogeneous networks (i.e., where neurons possess the same level of excitability), we harnessed spectral theory for large-scale random systems [59, 73]. By construction, our network model is subject to the circular law of random matrix theory [58, 73] in which the complex eigenvalues are constrained with high probability in a disk in the complex plane, with a spectral radius Γ centered around the local relaxation gain *d* (see MATERIALS AND METHODS). Changes in the spectral radius Γ result either in the clustering or dispersion of eigenvalues around the center of the spectral disk (i.e., d). As such, whenever the spectral radius Γ becomes larger (resp. smaller) than |*d*|, the network is said to become unstable (resp. stable) with high probability: eigenvalues cross the imaginary axis and exhibit positive (resp. negative) real parts [34, 36, 58, 74–76]. If the spectral radius remains commensurate with |*d*|, then the network is considered metastable and in the vicinity of a critical point. This framework has been used extensively in macroecology to examine the stability of complex natural systems, such as food webs [14, 16, 26–32].

While the net size of the spectral radius determines the system’s stability, how this spectral radius *changes* with respect to a control parameter (e.g., modulatory input amplitude *S_o_*) reflects the system’s resilience or volatility. That is, changes in spectral radius illustrate the system’s susceptibility to stability transitions due to changes in a control parameter. We thus subjected the homogeneous network to a thorough spectral analysis (cf. section 4.3). By virtue of having identical excitability, individual neurons’ steady states were found to be identical across the network and entirely dependent on the modulatory input amplitude (i.e. 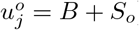), as expected. This is fully consistent with the dynamics observed in Fig. 2. Over short time scales, the modulatory input *S*(*t*) ≈ *S_o_* can be considered constant: its influence on the spectral radius Γ may thus be quantified. Indeed, as can be seen in Fig. 3A both numerically and analytically, the spectral radius was found be highly sensitive to modulatory input: changes in So resulted in high amplitude clustering and/or dispersion of the eigenvalues around the relaxation gain, causing frequent transitions between stability and instability. The spectral radius Γ was found to increase with the modulatory input amplitude (*S_o_*), indicating that such fluctuations generally lead to instability.

**Figure 3.**
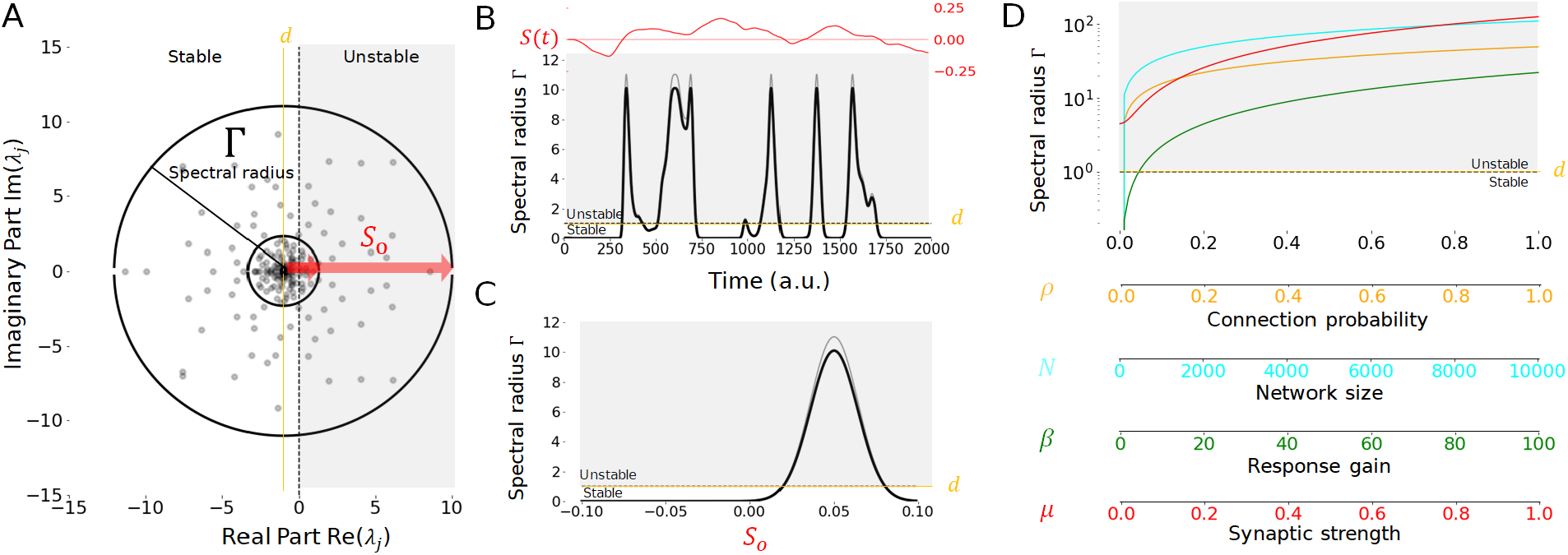
Spectral analysis of homogeneous networks exposed to modulatory input. Modulatory inputs influence the statistical properties and distributions of eigenvalues for homogeneous networks, which may be quantified by the spectral radius Γ. **A**. The network eigenvalues (λ_*j*_, computed for one instance of the network connectivity; gray dots) are complex and distributed in the complex plane within a disk (black circle), centered around the linear relaxation gain *d* (yellow vertical line) and delineated by a circle of radius Γ. Whenever Γ matches or exceeds the stability threshold (vertical black dashed line located at 0), the system is considered unstable (gray shaded area) with high probability. The slowly fluctuating modulatory input *S*(*t*) ≈ *S_o_* influences the system’s stability by expanding or contracting the spectral radius, and hence the spectral disk containing eigenvalues. As the modulatory input amplitude |*S_o_*| increases (horizontal red arrows) the spectral disk and radius increases, resulting in instability. Here, three examples are plotted for *S_o_* = 0 (small black circle), *S_o_* = 0.025 (medium black circle) and *S_o_* = 0.05 (large black circle). **B**. When *S*(*t*) fluctuates slowly in time (top red line), the spectral radius Γ expands and contracts above or below the stability threshold (Γ = *d* = 1; orange horizontal line) leading to alternating epochs of stability and instability (gray shaded area) as exemplified in Figure 2. Aside from changes in the amplitude *S_o_*, other parameters remained fixed. **C**. Spectral radius Γ as a function of *S_o_*. At baseline (i.e. *S_o_* = 0), the spectral radius is small and hence the network is stable. As |*S_o_*| increases, the spectral radius increases, exposing the system to stability transitions as eigenvalues cross the imaginary axis. As the modulatory input increases further, the spectral radius starts to decrease as the neurons reach saturation. The threshold of stability is plotted for Γ = |*d*| = 1 (see MATERIALS AND METHODS; orange horizontal line), alongside both numerically (grey) and theoretically (black) computed spectral radius Γ. Instability region is shaded in gray. **D**. Changes in connection probability (*ρ*; orange line), network size (*N*; cyan line), firing rate response gain (*β*; green line) and mean synaptic strength (*μ*; red line) are all collectively destabilizing and increase monotonically the spectral radius Γ. In this panel, *S_o_* = |*B*|. Each parameter was varied independently within the range specified, while other parameters were set to their default value i.e. *N* = 100, *ρ* = 0.05, *d* = −1, *μ_e_* = *μ* = 0.08, *β* = 50, *f* = 0.8, *μ_i_* = *fμ*/(*f* – 1), 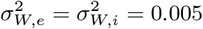, *B* = −0.05.

We confirmed this volatility in Fig. 3B, alongside the alignment between our numerical and analytical calculations. Time-dependent changes in the amplitude of the modulatory input (such as those exemplified in Fig 2) significantly contract and/or expand spectral radius Γ, whose value intermittently crosses the stability threshold, leading to an alternation between stability and instability. As *S_o_* fluctuates, the network undergoes epochs of instability, alternating with periods where neural activity is either suppressed (*S_o_* strongly inhibiting) and/or saturated (*S_o_* strongly exciting). We note that fast changes in *S*(*t*) might cause the network to cross the unstable regime briefly; instability is then difficult to observe since the system does not evolve sufficiently fast to exhibit unstable observable dynamics. Results plotted in Fig. 3C show a high dependence of the spectral radius on modulatory input amplitude (*S_o_*). Stability (i.e., relaxing neural activity, small Γ) characterizes inhibitory and/or low amplitude modulatory input, while higher amplitudes lead to instability (i.e., divergent, chaotic and/or synchronous neural activity, large Γ) and eventually saturation (i.e., neural activity plateaus, small Γ).

What must be concluded from these observations is that the spectral radius size is a context- and modulation-dependent metric for stability. Indeed, as neural systems can reside in both stable (relaxation) and/or unstable (oscillations, synchrony, chaos) functionally meaningful dynamic regimes, the spectral radius evaluated at a given moment in time conveys little information about the network dynamic volatility and resilience. It is instead how it *changes* that reflects resilience or volatility. As shown in Fig. 3D, our analysis also revealed that the spectral radius Γ - and hence the dispersion of eigenvalues in the complex plane - increases with network size (*N*), connection probability (*ρ*), firing rate response gain (*β*) as well as net synaptic strength (*μ*); individually or collectively, all these network features diminish the system’s resilience. This is in line with previous results [28, 34] notably on balanced networks [68, 69], highlighting that homogeneous networks are generically prone to instability. Taken together, our analysis indicates that, in sparse balanced and homogeneous networks, the spectral radius’ high sensitivity to modulatory input and other control parameters underlies the system’s changing stability, and thus its volatility.

### 2.3 Intrinsic excitability heterogeneity tunes stability and resilience

Numerous previous studies [45, 47, 57, 72] have shown that heterogeneous neural systems adapt and converge towards a regime of metastability to optimize responses and coding properties. Such metastability manifests itself through critical-like neural activity [77–79] and/or dynamics residing in the vicinity of a state transition [80]. From the perspective of the aforementioned circular law, such dynamical properties emerge whenever these networks are brought towards and operate in dynamical regimes resulting from a spectral disk of intermediate size, neither too small (i.e., strong stability leading to quiescence) nor too large (i.e., strong instability leading to divergence, chaos and/or synchrony).

Our previous findings [43] suggest that excitability heterogeneity should improve network resilience, as does the result presented in Fig. 1B. To further explore this we first repeated the numerical experiment in Fig. 2 in which network response to slow-varying modulatory input is examined over long time scales, but now in presence of excitability heterogeneity (i.e., 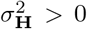). We exposed the network to the same connectivity statistics and modulatory input as before, while examining the difference in its behavior. In contrast to the homogeneous case, the long term dynamics of the network were found to be resilient: robust, invariant stability replaced the intermittent behavior seen in the homogeneous case. As can be seen for the simulations in Fig. 4A, no transitions between stability and/or instability occurred, and neuronal responses were qualitatively similar, smoothly driven by modulatory input amplitude. Neural activity remained in a regime in which perturbations relax back to equilibrium. As a direct consequence of heterogeneity, degeneracy in the neurons’ equilibria is broken: neuron fixed points were now distributed with a mean 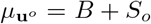 and variance 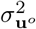 (Fig. 4C; see MATERIALS AND METHODS). Lyapunov exponents remained bounded below zero throughout, as can be seen in Fig. 4B. This behavior was also found to persist over independent realizations of the network connectivity (Fig. 4D). Take together, these confirm persistent stability and suggest enhanced resilience in presence of excitability heterogeneity.

**Figure 4.**
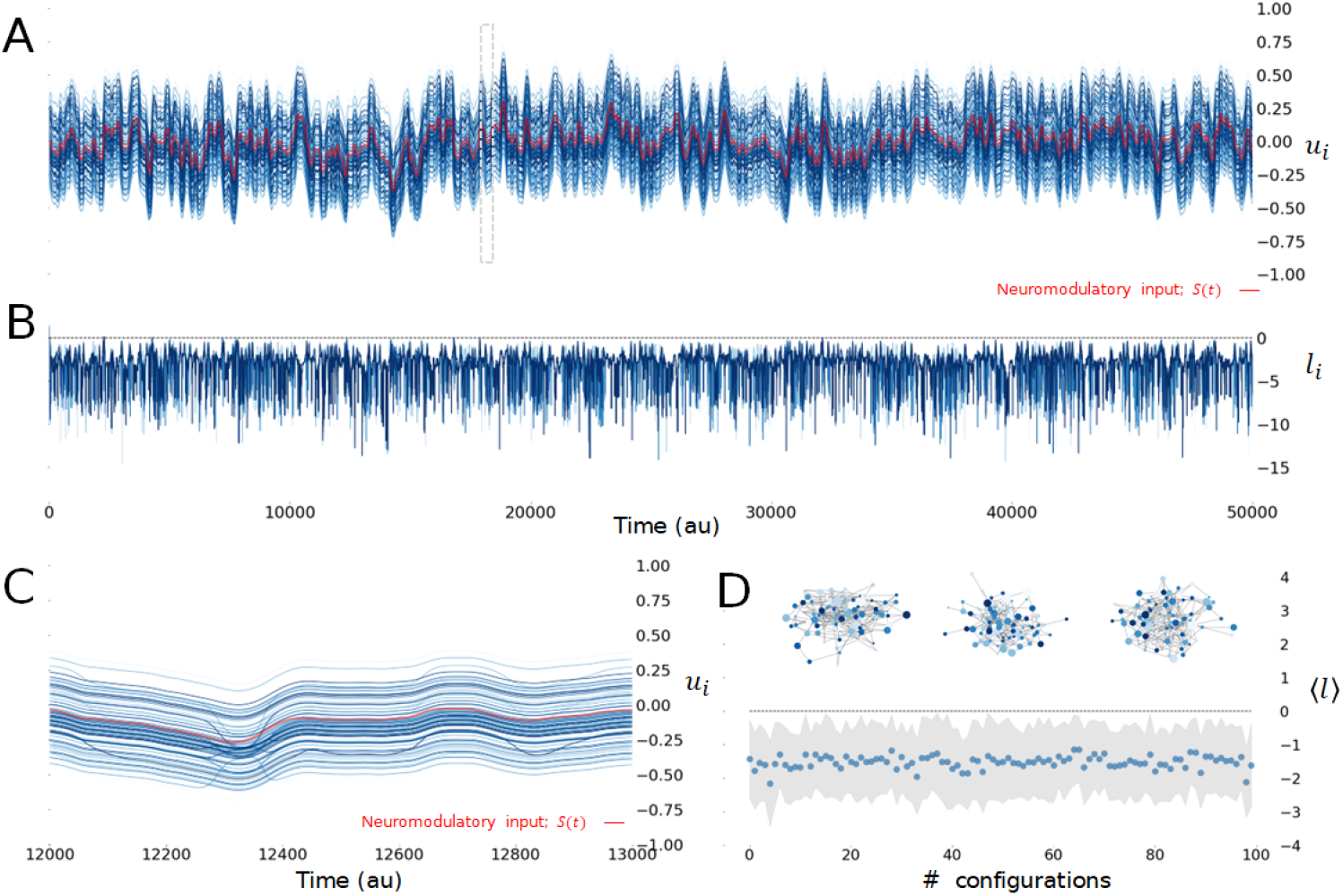
Heterogeneity quenches volatility and promotes resilience to modulatory drive over long time scales. **A**. Long time scale dynamics of an heterogeneous network 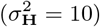 exposed to the same slowly varying drive (*S*(*t*); red line) as in Fig. 2. Individual neuron activity (*u_i_*; blue shaded lines) are now distributed around a mean activity 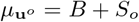 and variance 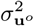. The network stability is preserved throughout: neuronal activity sits in a regime where it relaxes back to equilibrium under perturbations, and supervene to modulatory input. **B**. Lyapunov exponents (*l_i_*) – computed numerically based on time series of each neuron across time. Compared to the homogeneous case, stability persists as the Lyapunov exponents remain negative throughout. **C**. For all epochs, the network is stable and exhibits dynamics smoothly driven by the modulatory input. **D**. Network mean Lyapunov exponent 〈*l*〉 over independent network configurations, realized using identical parameters but different net connectivity. Fluctuations below the horizontal line (〈*l*〉 = 0; black dashed line) indicate resilient dynamics, in which no stability transition occurs. Gray shading indicates ± SD computed over independent trials of duration *T* = 3000a.u..Parameters are *N* = 100, *ρ* = 0.05, *β* = 50, *d* = −1, *f* = 0.8, *μ_e_* = *μ* = 0.08 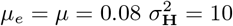, *μ_i_* = *fμ*/(*f* – 1), 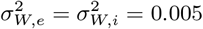, and *B* = −0.05.

To better understand the mechanism behind these dynamics, we adapted the spectral theory for large-scale random systems [59,73] to expose the influence of excitability heterogeneity on the distribution of eigenvalues. We specifically explored the susceptibility of the spectral radius Γ - and hence the dispersion of eigenvalues in the complex plane - to modulation across various degrees of heterogeneity 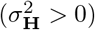 (see MATERIALS AND METHODS). Our analysis revealed two main roles played by diversity on network dynamics: a) homeostatic control on network stability; and b) the promotion of its resilience.

Indeed, we found that excitability heterogeneity is a homeostatic mechanism exerting bidirectional and context-dependent control on network stability: enriching the dynamics whenever they are too poor, or conversely, stabilizing network activity whenever it is too unstable. Indeed, as shown in Fig. 5A, heterogeneity increased the spectral radius (Γ) for small values modulatory input amplitudes (*S_o_*). For such low amplitudes of modulation, lack of heterogeneity yields highly stable neural activity that invariably relaxes back to equilibrium whenever perturbed: the spectral radius is infinitesimal and eigenvalues are clustered around the relaxation gain *d*. Introducing excitability heterogeneity expanded the spectral disk, enriching network dynamics towards instability. Surprisingly, higher modulatory input amplitudes, for which the system is highly unstable, led to the opposite. Indeed, heterogeneity was found to here instead contract the spectral disk and stabilize the dynamics (Fig. 5B). This contextual control of excitability heterogeneity on stability which depends on modulatory fluctuations (cf. Figs. 5A, B) suggests that heterogeniety tunes the spectral disk - and hence eigenvalue dispersion - towards an optimal intermediate size.

**Figure 5.**
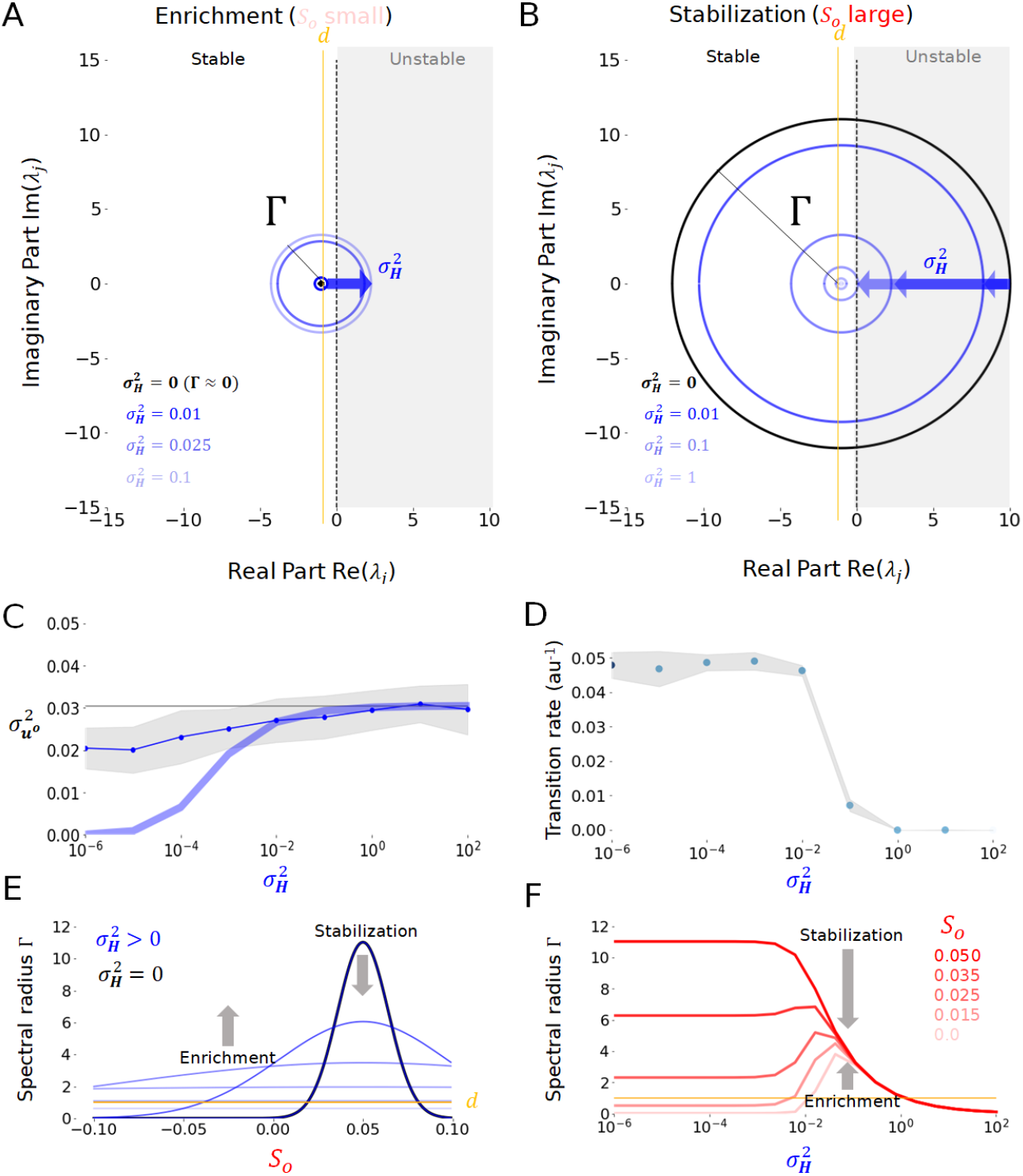
Heterogeneity-induced homeostatic control on stability. Increasing the degree of heterogeneity in the network strongly influences the network’s response to modulatory input. **A**. When modulatory input amplitude (*S_o_*) is small, increased diversity results in an enrichment of neural activity. As heterogeneity increases, the spectral disk (Γ) expands (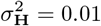, 0.025, 1; blue shaded circles) compared to the homogeneous case (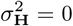; black circle). Modulatory input is here *S_o_* = 0. **B**. In contrast, when modulatory input amplitude is large, heterogeneity stabilizes neural activity by contracting the spectral disk compared to the homogeneous case (circle colors as in Panel A), causing a clustering of the eigenvalues around the linear relaxation gain *d* (vertical yellow line). Modulatory input is here *S_o_* = |*B*|. **C** Variance of the steady state distribution 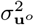 as a function of heterogeneity 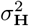. As heterogeneity increases, the variance of the steady state distribution increases. The dotted line corresponds to the numerically computed steady state distribution variance averaged over 50 independent network realizations. The bold blue line represents the analytical calculations in which the approximation 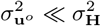 was used. Error bars reflect standard deviations over trials. **D** Stability transition rate as a function of excitability heterogeneity. This rate corresponds to the number of bifurcations per unit time over independent realizations of the network, for 10 trials of duration 50 a.u.. Error bars reflect standard deviations over trials. **E**. Spectral radius Γ as a function of excitability heterogeneity (colors as in Panel A). Diversity has an enrichment effect for low modulatory input, while being stabilizing whenever modulatory input is strong and/or saturating. **F**. The homeostatic influence of heterogeneity on the spectral radius depends on modulatory input. Diversity will invariably stabilize the network (i.e. decrease Γ) whenever *S_o_* is high (bold red curve), while enrichment (i.e. increased Γ) will occur for weak *S_o_* (pale red curve). High levels of heterogeneity are always stabilizing as Γ decreases to zero. Other parameters are given by *N* = 100, *ρ* = 0.05, *d* = −1, *f* = 0.8, *μ_e_* = *μ* = 0.08, *β* = 15, *μ_i_* = *fμ*/(*f* − 1), 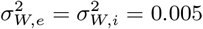 and *B* = −0.05.

To confirm the alignment of our mathematical analysis and numerical simulations, we computed the variance of the neuron’s fixed point distribution (i.e., 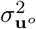), which was also found to depend on the degree of heterogeneity (Fig. 5C). Introducing heterogeneity also consistently prevented stability transitions, rendering the system more resilient. Indeed, as can be seen Fig. 5D, the transition rate - corresponding to the number of bifurcations observed in the network per unit time - decreased monotonically, confirming the trend seen in Fig. 4. We systematically quantified how excitability heterogeneity shapes the spectral radius in presence of modulatory fluctuations. As plotted in Fig. 5E, the contextual influence of excitability heterogeneity on network stability stems from a damping of spectral radius sensitivity with respect to modulatory input. Indeed, sharp changes in Γ caused by *S_o_* (such as those seen in Fig. 3B, C) were evened out by heterogeneity, resulting in an enrichment or stabilization of the dynamics as the spectral radius is increased or decreased, respectively. Specifically, heterogeneity increased the spectral radius for low and/or saturating modulatory amplitudes, and did the opposite for high amplitudes and decreased the spectral radius. The homeostatic influence of heterogeneity on network stability could be confirmed in Fig. 5F. Irrespective of modulatory input amplitude *S_o_*, heterogeneity was found to tune the spectral radius - through either enrichment or stabilization - towards the same intermediate radius.

Another important conclusion stemming from our analysis is that excitability heterogeneity generically enhances network resilience. As can be seen from Fig. 6A, increasing excitability heterogeneity significantly damped spectral radius changes resulting from modulatory input. Indeed, excitability heterogeneity made Γ less sensitive to changes in *S_o_*, and by doing do, quenched volatility. This was confirmed in Fig. 6B by systematically varying modulatory input amplitude and the degree of heterogeneity while measuring the spectral radius. We found that heterogeneity damped the sensitivity of the network stability on *S_o_*, as the spectral radius gradually radius becomes effectively independent of *S_o_* beyond a given degree of heterogeneity (dashed box in Fig. 6B). This implies that excitability heterogeneity anchors eigenvalue distributions in the complex plane, while making eigenvalues independent of modulatory input. An important consequence of this anchoring is that the network stability remains fixed, preventing stability transitions, confirming resilience. To encapsulate the effect of excitability heterogeneity on the network’s resilience, we computed both the spectral volatility (*κ*)- which measures the effective sensitivity of the spectral radius on a given control parameter - as well as the resilience parameter 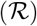 - which is the reciprocal of the spectral volatility - as a function of modulatory input amplitude (i.e. *S_o_*). These metrics quantify how invariant to changes in a given control parameter the eigenvalue distribution is. This is done by looking at variations of the spectral radius, cf. section 4.5. As shown in Fig. 6C, heterogeneity optimized resilience to modulatory input, and the spectral volatility decreased. Collectively, these results demonstrate that excitability heterogeneity, greatly enhances the resilience of sparse balanced networks by anchoring the eigenvalues in the complex plane and decreasing the sensitivity of their distribution to modulatory input.

**Figure 6.**
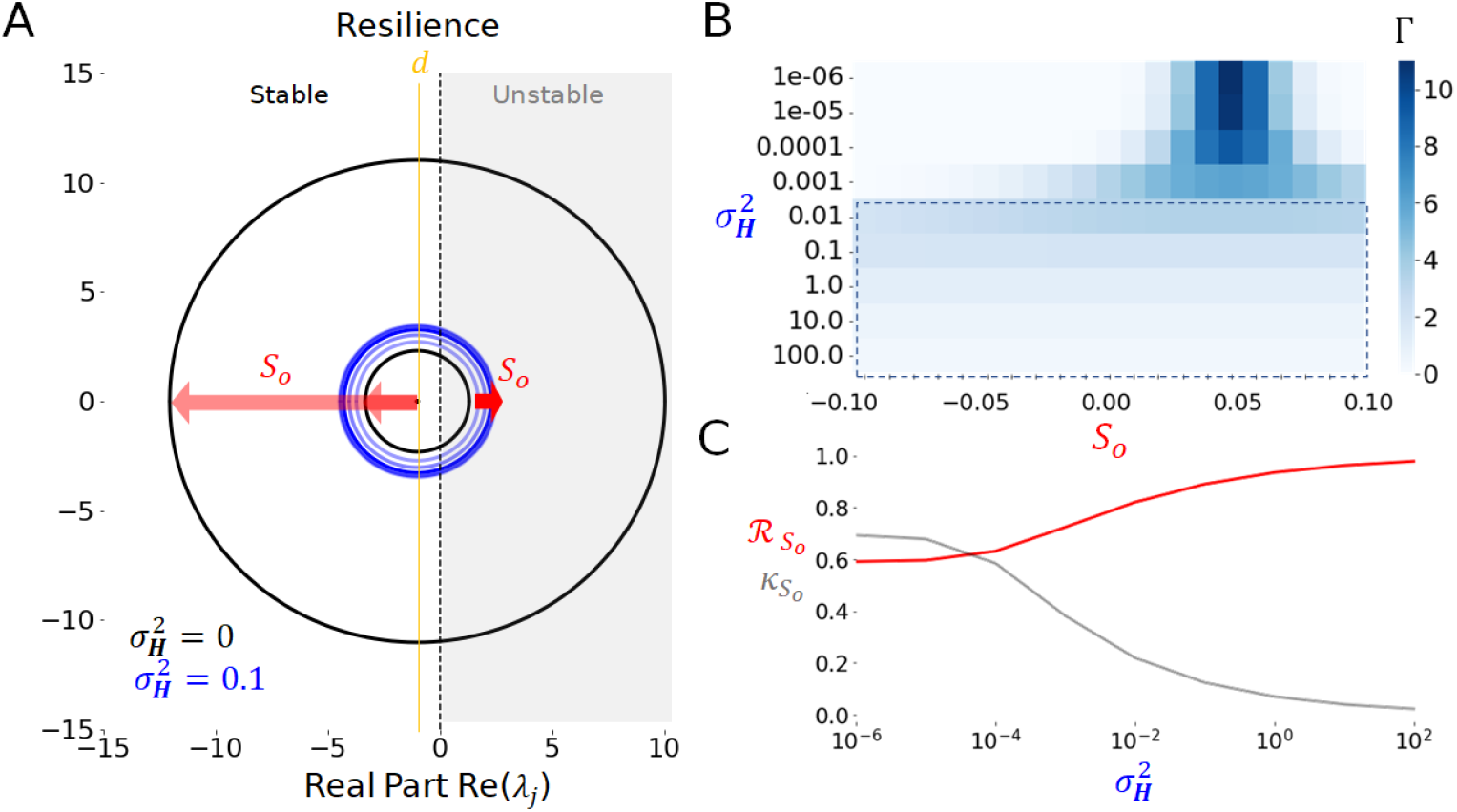
Heterogeneity enhances the resilience of balanced networks. **A**. Diversity limits variations of the spectral disk. Increasing heterogeneity (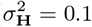; blue shaded circles) constrains changes in the spectral radius Γ resulting from changes in modulatory input amplitude (*S_o_* = 0, 0.025 and 0.05; red arrows; right). Without heterogeneity, the same fluctuations in modulatory input (red arrows; left) result in much wider changes in Γ (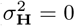, black circles). While both heterogeneous and homogeneous cases result in an expansion of the spectral disk, these variations are much smaller whenever 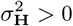; **B**. Spectral radius (Γ) as a function of the degree of heterogeneity and the amplitude of the modulatory input (*S_o_*). Increasing heterogeneity (dashed box) suppresses the system’s dependence on the modulatory input as the spectral radius Γ becomes constant despite changes in *S_o_* (dashed box). **C**. Resilience (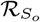; red curve) and spectral volatility (*κ_S_o__*; grey curve) measures with respect to the modulatory input amplitude (*S_o_*) as a function of the degree of heterogeneity. Resilience increases with heterogeneity while the spectral radius sensitivity (i.e. volatility) decreases with 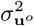. Other parameters are given by *N* = 100, *ρ* = 0.05, *d* = −1, *f* = 0.8, *μ_e_* = *μ* = 0.08, *β* = 15, *μ_i_* = *fμ*/(*f* − 1), 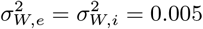 and *B* = −0.05.

### 2.4 Heterogeneity may stabilize networks across changes in connectivity

Our results so far indicate that excitability heterogeneity implements a long time-scale homeostatic control mechanism that promotes resilience in networks exposed to modulatory inputs. However, other control parameters might influence the neural systems’ stability over these time scales. Neural systems are subjected to perpetual change, even in the absence of modulatory fluctuations and/or stimuli. Synaptic plasticity is a salient example: during learning, the number of synapses and/or the effective synaptic weights increase, as a consequence of processes such as long-term potentiation (LTP) and depression (LTD) [12]. Networks undergoing such plasticity-induced structural modifications of their connectivity tend to be weakly resilient. Indeed, most forms of synaptic plasticity lead to the development of instability, in which run-away neural activity departs from baseline and needs to be compensated/stabilized through various homeostatic feedback processes [5, 63, 64], a few of which have found experimental support [81].

We asked whether excitability heterogeneity, on its own, could prevent stability transitions in neural systems undergoing plasticity-induced changes in connectivity. Our previous analysis demonstrates that the spectral radius Γ - and hence the dispersion of eigenvalues in the complex plane - increases with connection probability (*ρ*) as well as net synaptic strength (*μ*). This suggests that long time-scale changes in these control parameters - prone to increase together or independently during learning - generically promote instability and volatility. As can be seen in Fig.7, this is confirmed numerically: increasing both the connection probability (Fig.7A) and synaptic strength (Fig.7B) over a physiologically realistic range resulted in instability, as measured with the mean Lyapunov exponent. However, this only occurred in the homogeneous case 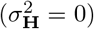. Indeed, increasing the heterogeneity suppressed this instability with the mean Lyapunov exponent remaining negative over the range of values of explored, i.e. the system did not experience any stability transitions.

**Figure 7.**
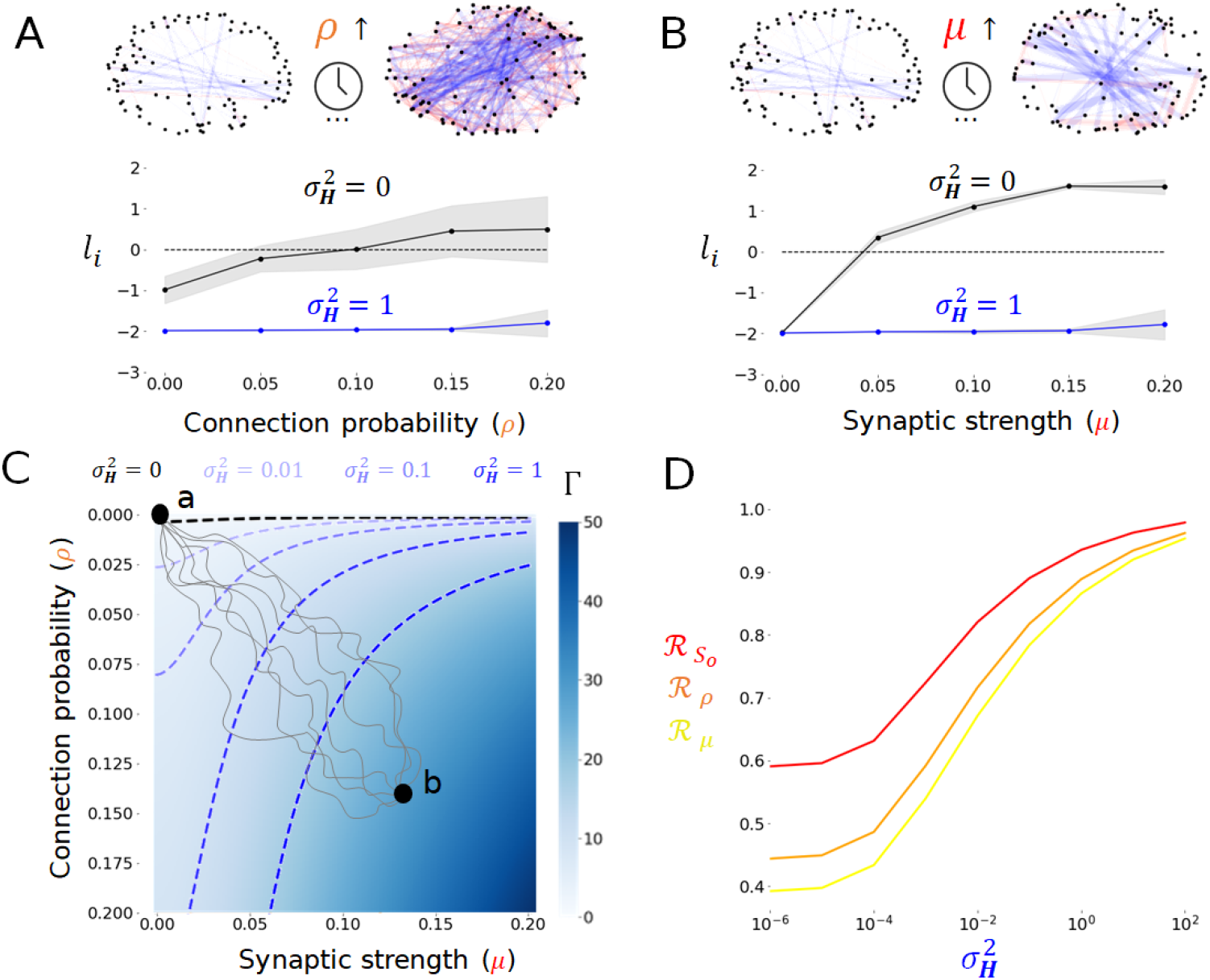
Heterogeneity compensates the destabilizing effect of changes in connection probability and synaptic strength. **A**. Increasing the connection probability generically destabilizes sparse balanced networks. In the homogeneous case (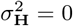; black curve), increasing the connection probability from *ρ* = 0 to 0.2 leads the network into an unstable regime. Lyapunov exponents *l_i_* increase and become positive. In contrast, in the presence of heterogeneity (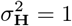; blue curve), they remain more or less constant, and stability persists. Here *μ_e_* = *μ* = 0.08, *μ_i_* = *fμ* (*f* − 1) **B**. The same trend is observed whenever the synaptic strength is increased from *μ* = 0 to *μ* = 0.2. Here *ρ* = 0.05. **C**. The spectral radius Γ increases with both connection probability (*ρ*) and synaptic strength (*μ*), suggestive of instability. The instability threshold (Γ(*ρ*, *μ*) = |*d*|) is shown as a black dashed line. Introducing excitability heterogeneity shifts the instability threshold in parameter space (blue shaded dashed lines), promoting stability. Illustrative curves representing a trajectory in parameter space occuring during plasticity(grey curves), connecting the network state before (*ρ* = 0, *μ* = 0; **a**) and after learning (*ρ* > 0, *μ* > 0;**b**). **D**. Resilience measure, computed as a function of connection probability (*ρ*; 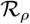; orange curve) and synaptic strength (*μ*; 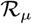; yellow curve), shown along modulatory input amplitude (*S_o_*; 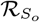; red curve) for reference. All these increase with increaseing degree of heterogeneity. Other parameters are given by *N* = 100, *d* = −1, *β* = 50, *f* = 0.8, 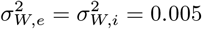. In panel D, *B* = 0.

Mathematical analysis affirms this finding. Figure7C shows that the spectral radius Γ increases monotonically with connection probability (*ρ*) and synaptic strength (*μ*) interchangeably, underlying such systems’ volatility and associated vulnerability to stability transitions. Slow time scale changes in connectivity resulting from plasticity (illustrated by the gray curves linking points **a** and **b** in Fig.7C) result in stability transitions. Introducing heterogeneity moved the effective stability threshold (i.e., Γ(*ρ*, *μ*) = |*d*|) further in parameter space, resulting in overall compensation for the destabilizing influence of increases in connection probability and synaptic strength (c.f., 3D). In this case, slow time scale changes in connectivity cause stability transitions to become increasingly unlikely as the net size of the stability region increases. In addition to this stabilizing influence, heterogeneity was also found to promote resilience by enhancing the persistence of stability via anchoring the eigenvalue distribution/spectral disk in the complex plane. We thus computed the resilience metric, now as a function of connection probability (*ρ*; 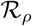) and synaptic strength (*μ*; 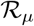). As shown in Fig.7D, increasing the degree of excitability heterogeneity enhanced resilience for both these control parameters i.e., promoting the persistence of stability by decreasing the spectral volatility and the susceptibility of the spectral radius on changes in connection probability (*ρ*) and synaptic strength (*μ*).

## 3 DISCUSSION

In the last several years, with continued advancements in high throughput [82] single cell RNA sequencing (scRNAseq) [83], and with the very recent addition of spatially resolved scRNAseq [84], it is abundantly clear that within cell-types there is a transcriptomic continuum rather than discrete sub-types [56]. This within cell-type transcriptomic diversity is also reflected in functional diversity in excitability features in human [4, 43, 85, 86] and rodent neurons [53, 56, 84] and likely a direct manifestation of the observed transcriptomic variability, given the correlation between the transcriptome and electrophysiological properties of neurons [87, 88]. In light of these technical advances in describing the properties of individual neurons at scale, a major challenge for neuroscience is to bridge across the divide between individual neuronal properties and network function [89]. While bridging this gap remains a significant challenge experimentally, although advances in imaging technologies (NeuroPixels [90], Ca2+ [91], ultrasound [92]) are continually closing it, it is the promise of computational and mathematical analyses to simplify the complexity of the brain while addressing this critical divide between brain structure and function [93].

It is within this context of bridging scales that we here bridge between neuronal diversity - a seemingly fundamental design principle of the brain - and the stability of cortical dynamics. We have been in part biased by our initial work in the context of epilepsy which is a pathological condition where individuals slip in and out of pathological dynamical brain states [24, 94] called seizures, and how excitability homogenization renders circuits more prone to such seizure-like states [43]. However, we have also have been greatly influenced by computational and mathematical work in macroecology that argue that not all types of diversity are stabilizing. Indeed, there is decades of research in the fields of macroecology and food webs, examining the relationship between complex systems’ stability, biodiversity and resilience: the so-called ”stability-diversity” debate [14, 16, 30, 33]. Fascinatingly, large scale random networks are more prone to volatility and stability transitions in response to increased size [14, 32, 34], connection probability [14, 31, 34–36], and connection strength [14, 26–28, 31, 35], and/or when connectivity motifs become too heterogeneous [26–28, 31, 32, 34–37].

Thus, within this stability-diversity debate and the myriad of heterogenities that could be explored, our choice to explore excitability heterogeneity is not haphazard, and for four reasons it is not surprising that we find that it has profound effects on resiliency of brain circuits. Firstly as discussed above, cellular diversity is the norm in the brain, and thus appears to be a clear “design principle” of neuronal circuits, which we accept at face value to be beneficial to the brain, and for which the biological machinery clearly exists [84, 95]. Secondly, there is ample evidence both experimentally and computationally that excitability heterogeneity is helpful for information coding in the brain, decorrelating brain networks while expanding their informational content [3, 71]. Thirdly, we have shown that amongst a number of experimentally determined electrophysiolgical features of human neurons, it is the loss of excitability heterogeneity that accompanies epilepsy [43]. Our mathematical and computational work showed that excitability heterogeneity prevents sudden transitions to highly correlated information poor brain activity. Lastly, neuronal excitability is highly malleable. This malleability arises from the process of *intrinsic plasticity*, where neuronal excitability is modulated by the neuron’s past activity [5, 61]. Indeed learning is accompanied by changes in voltage and calcium activated channels that are principally involved in setting resting membrane potential, input resistance, and rheobase [61]. It is these kinds of channels as well that are altered in a number of neuropsychiatric conditions, including epilepsy [62]. Excitability thus represents a local parameter tuned to the complexities, or lack thereof of activity of each neuron in the sea of activity it is embedded in.

Furthermore, in light of the ubiquity of various forms of neuronal [4, 40, 96–99] and glial [41, 42] diversity, that could render neural circuits unstable (i.e., the stability-diversity debate above), they are in fact highly resilient, and qualitatively invariant across extended time scales in part likely due to excitability heterogeneity. This of course holds true in healthy brains despite continuous external and internal changes, driven by factors including modulatory inputs [6–11], environmental fluctuations and/or stimuli [1, 25, 60], and changes in connectivity like those resulting from synaptic plasticity [63, 64]. The robustness of neural dynamics and function - the persistence of its dynamics - with respect to changing control parameters epitomizes resilience [13–17, 30].

This also holds true for process that continuously change the brain during development and ageing, where brain dynamics remain stable over many decades despite the structural changes that accompany time, and pathological processes, where failure to regulate brain activity in the face of pathological insults predispose the brain to dynamic volatility [18, 21]. A confluence of both experimental [1, 4, 25, 38–43] and theoretical studies [43–50] have highlighted the role of heterogeneity in brain dynamics and stability. Notably, phenotypic diversity has been shown to promote the stability of brain function and its associated dynamics through degeneracy, redundancy and covariation [1, 25, 100].

Our initial results confirmed [68, 69, 101] that networks with homogeneous excitability exhibit volatility in response to modulatory input. Increases in network size, synaptic strength, and connection probability all led to stability transitions. These observations highlight that the spectral radius size itself, evaluated at a given moment in time, conveys limited information about a network’s volatility and susceptibility to critical transitions, which remains high in the absence of phenotype diversity. Introducing excitability heterogeneity changed the portrait completely: our joint numerical and analytical results revealed that excitability heterogeneity: 1) implements a homeostatic control mechanism tuning the distribution of eigenvalues in the complex plane in a context-dependent way; and 2) enhances network resilience by fixing this distribution and making it independent of modulatory input(s). We extended our analysis to connection probability and synaptic strength, parameters that are prone to change over long time scales during processes such as plasticity [63, 64] and development [102]. Excitability heterogeneity also promoted resilience here by preserving network stability.

This formalism also facilitates insightful observations about the role played by various forms of diversity in neural circuits, and complex natural systems generally. In [37], the authors provided a comprehensive overview of the destabilizing influence of motif heterogeneity - the variability in the connection degree, or alternatively a lack of redundancy in connectivity - on complex random graphs. Recontextualized from the perspective of neural systems, these results and ours suggest that networks exhibiting redundant connectivity motifs alongside node heterogeneity will generically exhibit enhanced stability and resilience, corroborating numerous experimental findings [1].

As a corollary to our results, less heterogeneous systems should be more vulnerable [18, 43] to critical transitions [18]. Epilepsy is a revealing example in which seizures, which are transitory events typified by hyper-active and -synchronous brain activity [103], seemingly occur paroxysmally. The occurrence of such seizures has been shown to depend on modulatory factors, such as stimuli [20] and circadian and/or multidien cycles [21]. While asynchronous activity relies on a controlled balance between excitation and inhibition [104], the transition to pathologically synchronous activity in epilepsy has largely been conceptualized as a disruption of this balance [19]. Our recent study [43] adds a new dimension to this mechanistic conceptualization of epilepsy, where we observed a decrease in excitability heterogeneity of layer 5 pyramidal neurons in seizure generating areas (epilepetogenic zone) in individuals with medically refractory epilepsy. When implemented computationally, this experimentally observed reduction in excitability heterogeneity rendered neural circuits prone to sudden dynamical transitions into synchronous states with increased firing activity, paralleling ictogenesis. These observations suggest an important contribution of neural heterogeneity - or lack thereof - in the disease’s etiologies.

Collectively, our results suggest that excitability heterogeneity make balanced sparse neural networks insensitive to changes in many key control parameters, quenching volatility preventing transitions in stability that typify a lack of resilience. This phenomenon is far from being exclusive to neural systems: the role of diversity in ecosystem resilience in the face of change has been extensively studied [14–16, 26–32], and extended across environmental science, ecology, engineering, operation research, management science, business, social sciences, and computer science [13, 14, 16, 17, 105]. The instrumental role of diversity at promoting resilience is reminiscent of the Gaussian blur effect observed in data analysis, in which high frequency gradients (i.e, “noise”) are filtered out to preserve the smoothness of data by suppressing non-linearity [106]. In the context of our work, such heterogeneous “blurring” occurs in control parameter space, resulting in a smooth (and eventually flat and decoupled) relationship between the spectral radius size and given control parameter(s). We thus argue that its not the spectral radius size itself, but how it changes in response to modulation, that reflects whether a system is resilient or not. Indeed, as neural systems (and complex natural systems generally) can reside in both stable (e.g., relaxation) and/or unstable (e.g., oscillations, synchrony, chaos) functionally meaningful dynamic regimes [18, 107], the spectral radius remains a context-dependent measure of stability that has little to do with the actual function of these systems. Dynamical invariance, robustness and resilience are consequences of the persistence of the spectral radius size over time and in the face of change, resulting from a heterogeneity-induced decoupling between spectral radius size and system’s control parameter(s) (e.g., modulation).

We highlight that our analyses and results can be generalized across a wide range of network sizes, connectivity profiles, topologies, types of heterogeneity, dynamics (e.g., asynchronous, rhythmic), and individual neuron response properties. Indeed, while we have focused here on Erdős–Rényi - type topology, our results may be easily extended to other graph structures (e.g., multi-modal, scale-free, cascade models) through a proper rescaling of the spectral radius [33, 37], and can also be modified to study time delayed systems [75]. In particular, the circular spectral disk resulting from the connectivity matrix considered here might adopt a different shape whenever predator-prey, competition and/or mutualistic interactions are introduced, yet are fully amenable to a node diversity considerations [60, 108].

Like all computational and theoretical work, there are limitations to the contexts in which these results are applicable. First, our model represents a balance between neurophysiological relevance and mathematical tractability. More detailed and biophysically rich models are certainly required to provide a more comprehensive understanding of the role of diversity on network stability. Second, phenotype diversity certainly impacts neural activity beyond excitability. A more thorough characterization of neural variability is surely warranted to reinforce the alignment between our model and experimental data, notably to improve the scope of our predictions. Third, our model does not consider stochastic fluctuations that are known to be ubiquitous in neural systems [109] and to influence their stability [110–112]. We have neglected this source of variation due to the long time scales considered here, and the moderate non-linearity of the system (i.e., parameterized by the firing rate response gain *β*). Future work is required to incorporate noise in both our simulations and analyses.

In summary, our results position excitability heterogeneity, and possibly more generally, neuronal diversity (from transcriptomic and functional studies), as a critical design feature of the brain to ensure its rich dynamics are preserved in the face of a wide set of network parameters. Furthermore, resilience of dynamics to scale (physical scale as in number of neurons, or connectivity as in the strength of connections) is of course critically important for a growing developing brain, as well as an ageing brain. However even more generally as a design principle, excitability heterogeneity provides dynamical resilience to brains of all sizes across the phylogenetic tree, allowing the *freeness* from scale in a scale-free system.

## 4 MATERIALS AND METHODS

### 4.1 Network model

We consider a large network of *N* neurons whose activity evolves according to the interplay between local relaxation, recurrent synaptic connectivity and slowly varying modulatory input. This model provides a description of dynamics unfolding over extended time scales, hence quantifying mean neuronal activity. The mean somatic membrane potential of neurons *u_i_*(*t*), *i* ∈ [1, *N*] obeys the following set of non-linear differential equations

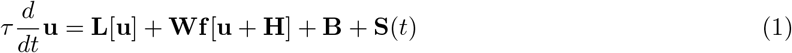

where *τ* scales the slow time scale at which the dynamics occur. In the following we re-scale time by *t* → *tτ* for convenience. The model in Eq. 1 is both flexible and general, encompassing the mean behavior of a wide scope of interconnected neurons models involving excitatory and inhibitory interactions, such as the celebrated Wilson-Cowan and Jansen-Rit models, for instance. Depending on the spatial scale considered, which remains here undefined, such models can be either be considered to be neuron-based (where nodes represent individual neurons - the perspective we adopt here) or populations (where nodes represent assemblies of such neurons), geared towards the characterization of neuronal mean activity across extended time scales. The term **L**[**u**] = *d***u** is a linear local relaxation term with rate *d* < 0 and 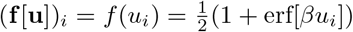 represents the firing rate response function of neurons with the Gaussian error function erf[**·**]. This function relates the membrane potential activity to the firing rate of the neuron. The vector-valued term **H** implements node diversity through spatially heterogeneous neuronal excitability, i.e. variable firing rate thresholds between neurons. The entries of **H** are sampled from a zero-mean Gaussian distribution of variance 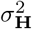. The neuron’s baseline activity **B** < **0** is a scaling factor used to set the neurons in a subthreshold regime in the absence of input. Lastly, the network in Eq. 1 is further subjected to a slow modulatory input **S**(*t*).

The connectivity matrix **W** in Eq. (1) specifies synaptic coupling between any pair of neurons. We assume randomly distributed excitatory and inhibitory coupling [76], with connection probability *ρ*. This connectivity motif corresponds to a weighted Erdős–Rényi random graph; we emphasize, however, that the following results may be easily extended to other topologies (e.g., [37]). The strength of these synaptic connections are individually Gaussian-distributed with mean *μ_e_* and *μ_i_*, variance 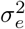 and 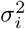 and with probability density functions *p_e_* and *p_i_*, respectively. We ensure that there are no self-connections i.e. *W_ii_* = 0 ∀*i*. In addition, we parametrize the relative density of excitatory versus inhibitory connections by a coefficient *f*, 0 ≤ *f* ≤ 1. Consequently, the probability density function of synaptic weights *W_ij_* may be written as *p* = *ρfp_e_* − *ρ*(1 − *f*)*p_i_*. Moreover, we choose a balanced connection connectivity with 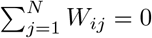, i.e. the sum over excitatory and inhibitory synaptic connections vanishes at each node. Given these constraints, the mean connectivity of the network is *μ_W_* ≡ E[*W_ij_*] = 0 and the variance of the synaptic connectivity becomes [76] 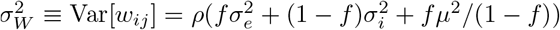. The mean network activity 〈*u*〉(*t*) is defined as the average activity across all neurons

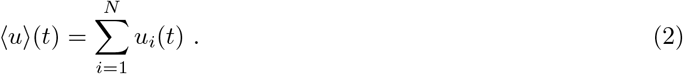

### 4.2 Stability

By construction, this network subscribes to the circular law of random matrix theory [58, 73]. According to this law, the statistical distribution of eigenvalues of the network - reflecting stability - is constrained with high probability within a disk centered around the local relaxation gain *d* (called the spectral disk) in the complex plane, whose radius Γ can be determined analytically. The eigenvalues populating that disk are complex numbers: if the disk is bounded in the left hand side of the imaginary axis (i.e., all real parts of these eigenvalues are negative), the network is said to be stable and its activity invariably relaxes back to its equilibrium after a perturbation. In our network model, such stable equilibrium is characterized by weak, asynchronous neuronal firing. If some eigenvalues cross the imaginary axis (i.e., the spectral disk is too large and some eigenvalues possess positive real parts), the network is said to be unstable, leading to activity that diverges, is synchronous and/or chaotic. In the intermediate case, when dominant eigenvalues (those possessing the largest real part) exhibit a near-zero real part, the network is said to reside at a critical point sitting between stability and instability, commonly referred to as metastable.

The stability of Eq. (1) may hence be characterized through the circular law [34, 36, 73]. Over a short time scale, modulatory input can be considered constant, i.e. **S**(*t*) = **S**, and the fixed point **u**^*o*^ satisfies

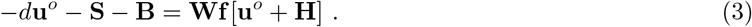

The stability of these fixed points can be determined by considering the spectrum Λ of the Jacobian matrix *J* of Eq. (1)

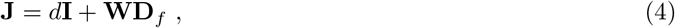

where **I** is the *N*-dimensional identity matrix and 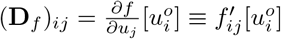 corresponds to the derivatives of the transfer function *f* evaluated at the fixed point **u**^*o*^.

It is well known [73] that for large *N* the spectrum Λ of Eq. (4) may be decomposed into an edge (Λ_*e*_) and bulk (Λ_*b*_) spectrum, i.e. Λ = Λ_*e*_ + Λ_*b*_. The real eigenvalues populating the edge spectrum λ_*e*_ ∈ Λ_*e*_ have here the mean λ_*e*_ = *d*. The circular law states [58, 73] that the remaining complex eigenvalues λ_*b*_ ∈ Λ_*b*_ populating the bulk spectra are confined within a disk of radius Γ given by

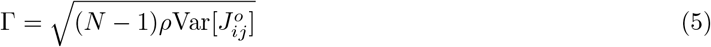

with 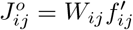 evaluated at the fixed point **u**^*o*^. This implies the maximum eigenvalue real part is given by max[*d*, *d* + Γ]. Since *d* < 0, the system’s stability is fully determined by the bulk spectrum and the fixed point **u**^*o*^ is stable if

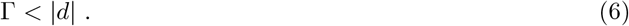

Used together, equations 5 and 6 are key to determining the influence of the network properties on its stability.

### 4.3 Stability of homogeneous networks

In absence of heterogeneity and modulatory input, i.e. **H** = 0 and **S** = 0, the fixed point **u**^*o*^ defined in Eq. (3) has the degenerate solution **u**^*o*^ = **B** = *B***1**^*T*^ with **1**^*T*^ = (1, 1 …, 1)^*T*^ and *B* < 0. Then elements of the Jacobian for **u**^*o*^ = **B** can be computed as

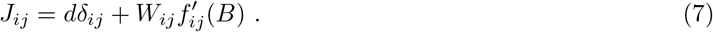

The spectral radius (5) can readily be determined in this case by computing the variance of the Jacobian as per Eq. 5 to

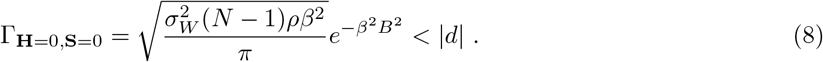

Whenever modulatory input is present, i.e. **S** = *S_o_***1**^*T*^ with **1**^*T*^ = (1, 1, .., 1)^*T*^, the fixed point becomes **u**^*o*^ = (*S_o_* + *B*)**1**^*T*^. For later purpose, we state in addition that the elements of the fixed point solution for **H** = **0** obeys a probability density function with mean *μ*_**u**^*o*^_ = *S_o_* + *B* and vanishing variance 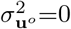. The stability criterion for **H** = **0**, **S** ≠ **0** now reads

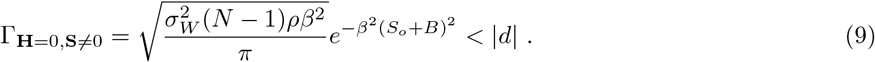

Equation (9) reveals that increasing the network size *N*, connection probability *ρ*, the variance 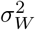 of synaptic weights (implying increases of the mean *μ* and excitatory and inhibitory variances *σ_e_*, *σ_i_*) as well as the firing rate response gain *β* cause an expansion of the spectral radius Γ, as does the modulatory input amplitude *S_o_*. The dependence of Eq. (9) on these various parameters is plotted in Fig. 3. Consequently, they all lead to instability of Eq. (1), in line with previous studies [37, 111].

### 4.4 Stability of heterogeneous networks

The heterogeneous case with modulatory input **H** ≠ 0, **S** ≠ **0** is more involved. The influence of excitability heterogeneity on stability may nonetheless be exposed by investigating how diversity in excitability thresholds, i.e. **H**, impacts the spectral radius Γ. Recall that **H** are random and sampled from a distribution *p*_H_ assumed to be Gaussian with zero mean and variance 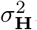. The fixed point **u**^*o*^ of Eq. (1) satisfies, instead of Eq. (3),

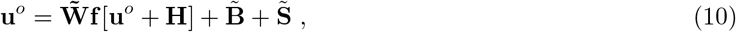

with 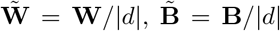 and 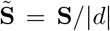. Equation (10) is a discrete version of the Hammerstein equation [113], whose solution is distributed with some probability distribution *p*_**u**^*o*^_. We highlight the important distinction to the homogeneous case (**H** = 0) in which the corresponding fixed point was degenerate.

Consistent with analysis steps in the previous section, stability is determined via the variance of the Jacobian as per Eq. 5. The matrix elements of the Jacobian for **H** ≠ 0, **S** ≠ 0 are given by

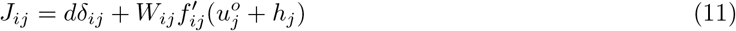

with *h_j_* = (**H**)_*j*_. Assuming independence between the fixed points **u**^*o*^ and the synaptic weights *W_ij_*

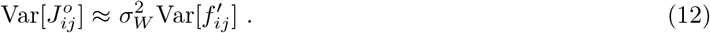

For large *N*, one may approximate this variance of the Jacobian by assuming that the distribution 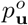 is independent from **H** and Gaussian-distributed with mean *μ*_**u**^*o*^_ and yet unknown variance 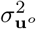. We find

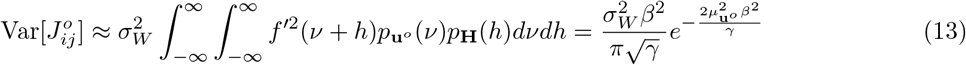

where 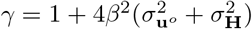. This result implies that stability depends on the mean and variance of the fixed point distribution *μ*_**u**^*o*^_ and 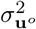, respectively, and the variance of the excitability threshold distribution 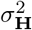. Since the excitability threshold variance can be chosen independently but determines the fixed point by Eq. (10), it is necessary to compute the mean *μ*_**u**^*o*^_ and variance 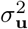 of the fixed point probability density function *p*_**u**^*o*^_.

At first, recall that homogeneous excitability, i.e. 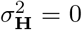 as described in section 4.3, implies 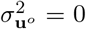. Consequently, 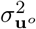 depends on 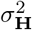 implicitly for heterogeneous excitability. In the following we assume modulatory input being homogeneous over the network. Then, in line with previous calculations

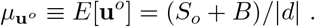

Moreover, with Eq. (10)

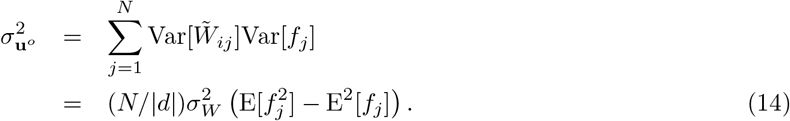

with 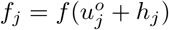. Then, we find for large *N*

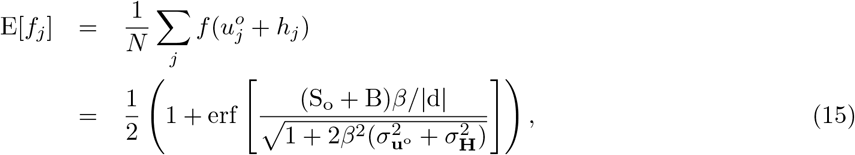

and

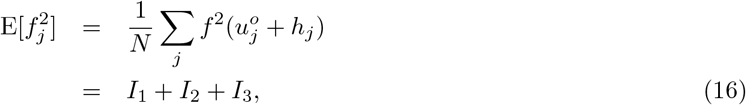

with *I*_1_ = 1/4, *I*_2_ = E[*f_j_*] − 1/2 and

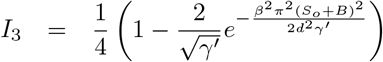

where 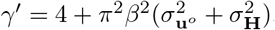. Here we have used the good approximation 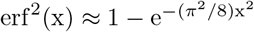.

Combining Eq. (14), (15) and (16) yields an implicit expression for the fixed point distribution variance

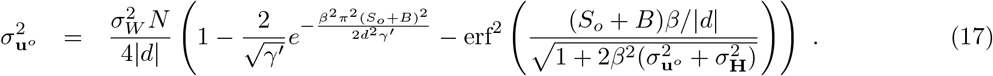

Equation (17) defines 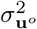 implicitly. In the absence of heterogeneities, i.e. 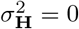, Eq. (10) stipulates **u**^0^ = 0 and 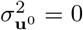 and Eq. (17) holds. Moreover, for excitability heterogeneities that are much stronger than fixed point fluctuations, i.e. 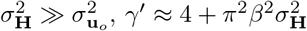 and Eq. (17) provides an explicit expression for 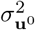.

With this result, the stability condition for the fixed point reads

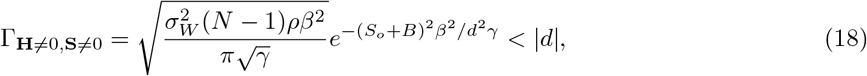

where 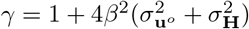, using the solution of Eq. (17).

### 4.5 Volatility and Resilience

Resilience refers to the qualitative invariance of dynamical states when exposed to changes in one control parameter, and the absence of stability transitions. To measure resilience and the robustness of eigenvalue distributions, one may quantify the sensitivity of the spectral radius Γ to changes in a given control parameter (i.e., how much the spectral radius fluctuates when exposed to changes in a certain parameter *P*).

We define the spectral volatility by

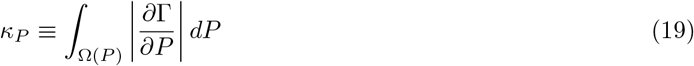

The spectral volatility *κ_P_* reflects the sensitivity of the eigenvalue distribution (the spectral disk area) to change in the parameter *P* over the range Ω(*P*) of values this parameter can take. It scales with how much Γ changes as a function of variations in the control parameter *P*: small volatility reflects persistence of the eigenvalue distributions and its overall resistance towards stability transitions. Specifically, if 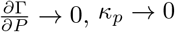.

One can use Eq. (19) to quantify the persistence of the eigenvalue distribution and spectral radius to changes in a control parameter. We thus introduce the resilience measure 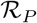 with respect to the control parameter *P* by considering the reciprocal of the spectral volatility

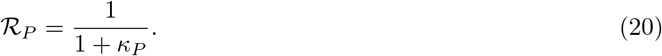

Note that whenever 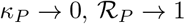 and if 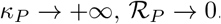.

Since we have derived the spectral radius analytically in Eq. (18), we are in a position to compute the spectral volatility and resilience as a function of all model parameters. Specifically, if one considers *P* = *S_o_*, for Ω(*S_o_*) = (−∞, +∞) one obtains,

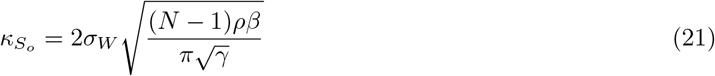

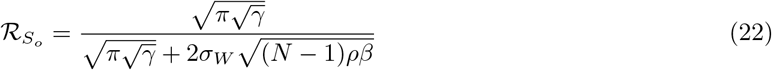

where 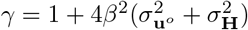. Equations (21) and (22) show that the volatility and resilience of the network with respect to the modulatory input amplitude both depend on excitability heterogeneity through the factor *γ*. Whenever 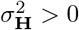 increases, the spectral volatility *κ_S_o__* decreases and the resilience 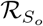 increases.

